# The nucleoporin ELYS regulates nuclear size by controlling NPC number and nuclear import capacity

**DOI:** 10.1101/510230

**Authors:** Predrag Jevtić, Andria C. Schibler, Gianluca Pegoraro, Tom Misteli, Daniel L. Levy

**Author notes:** Co-first authors (Predrag Jevtić, Andria C. Schibler). Corresponding authors: Daniel L. Levy, University of Wyoming, Department of Molecular Biology, 1000 E. University Avenue, Laramie, WY, 82071, Tom Misteli, National Cancer Institute, NIH, Center for Cancer Research, 41 Library Drive, Bldg. 41, B610, Bethesda, MD, 20892.

## Abstract

How intracellular organelles acquire their characteristic sizes is a fundamental cell biological question. Given the stereotypical changes in nuclear size in cancer, it is particularly important to understand the mechanisms that control nuclear size in human cells. Here we use a high-throughput imaging RNAi screen to identify and mechanistically characterize ELYS, a nucleoporin required for postmitotic nuclear pore complex (NPC) assembly, as a determinant of nuclear size in mammalian cells. We show that ELYS knockdown results in small nuclei, the accumulation of cytoplasmic lamin aggregates, reduced nuclear lamin B2 localization, lower NPC density, and decreased nuclear import. Increasing nuclear import by importin α overexpression rescues nuclear size and lamin B2 import, while inhibiting importin α/β nuclear import decreases nuclear size. Conversely, ELYS overexpression leads to increased nuclear size, enrichment of nuclear lamin B2 staining at the nuclear periphery, and elevated NPC density and nuclear import. Consistent with these observations, knockdown or inhibition of exportin 1 increases nuclear size. In summary, we identify ELYS and NPC density as novel positive effectors of mammalian nuclear size and propose that nuclear size is controlled by nuclear import capacity.

## INTRODUCTION

A fundamental question in cell biology is how intracellular organelles acquire their characteristic sizes. Organelle size almost certainly impacts function. For instance, expansion of the endoplasmic reticulum (ER) occurs in response to the need for increased protein synthesis or folding [1–3]. Depending on cell-size and -type, Golgi and mitochondrial volumes must be sufficient to support cellular metabolic requirements [4–8]. The functional significance of nuclear size is a particularly important problem. Changes in nuclear size are used to diagnose and stage many forms of cancer [9–12], yet it is unknown if these nuclear size changes contribute to the disease pathology or result from it. Addressing this question requires a comprehensive understanding of the mechanisms that control nuclear size in normal and cancerous mammalian cells.

The nuclear envelope (NE) is composed of an enclosed double lipid bilayer that is continuous with the ER on the exterior and lined on the inside by multiple proteins that constitute the nuclear lamina, most notably nuclear lamins [13]. Inserted into the NE are nuclear pore complexes (NPCs) that mediate nucleocytoplasmic transport of proteins and RNA [14–17]. The NPC is a large protein complex composed of multiple copies of ~30 different proteins termed nucleoporins (Nups), and distinct mechanisms are responsible for assembling NPCs into the NE after mitosis and for inserting NPCs into the NE during interphase [18,19]. Classical nuclear import is mediated by importin α/β karyopherins that transport cargos through the NPC and into the nucleoplasm where RanGTP induces cargo release through importin β binding. The three major nuclear lamins (lamin A/C, B1, and B2) are imported through this pathway. Conversely, exportins complexed with RanGTP mediate cargo export. While nuclear Ran is GTP-bound due to the chromatin localization of its guanine nucleotide exchange factor (RCC1), cytoplasmically localized RanGAP converts RanGTP to RanGDP. Cytoplasmic Ran is recycled back into the nucleus by its dedicated import factor, NTF2 [20–24].

Nuclear sizing mechanisms have been identified in a variety of model systems [11,25,26]. In yeast, nuclear size scales with cell size and blocking nuclear export leads to an increase in nuclear size [27–29]. Using *Xenopus* egg extracts it was shown that differences in the levels of importin α and NTF2 account for nuclear size differences in two different *Xenopus* species [30]. Over early *Xenopus* development, changes in cytoplasmic importin α levels and protein kinase C activity contribute to reductions in nuclear size [30–32]. In *C. elegans*, nuclear size is sensitive to the levels of importin α, NTF2, and RCC1 [33,34]. Altering lamin expression levels impacts nuclear size in *Xenopus, C. elegans*, and mammalian cells [34–36]. Cytoskeletal elements also affect nuclear size. In confined *Xenopus* egg extracts, nuclear size scales with the size of microtubule asters [37]. In mammalian cell culture, nuclear filamentous actin promotes nuclear growth [38], while connections between cytoplasmic actin and nesprins in the outer nuclear membrane tend to restrict nuclear growth [39,40]. Because nuclear and ER membranes are contiguous, changes in ER morphology can also impact nuclear size [41,42].

While a yeast screen has been performed to identify nuclear size effectors [28] and model systems such as *Xenopus* and *C. elegans* have begun to reveal some conserved mechanisms of nuclear size regulation [43–45], questions remain about how nuclear size is regulated in human cells. Beyond testing known mechanisms of nuclear size regulation in mammalian cells, imaging-based RNAi screens offer an opportunity to identify novel nuclear size effectors [46]. We have performed a high-throughput imaging RNAi screen for nuclear size effectors in breast epithelial cells and here describe our mechanistic analysis of one candidate identified in the screen, ELYS (also known as MEL-28 and AHCTF1), one of the first Nups recruited to chromatin for postmitotic NPC assembly [47–51]. Previous work demonstrated that nuclei assembled in *X. laevis* egg extract failed to assemble NPCs when ELYS was immunodepleted or upon addition of a dominant negative fragment of ELYS and, as expected for import-deficient nuclei, no nuclear growth was observed [52,53]. Here we demonstrate that NPC densities are sensitive to ELYS protein levels in cultured mammalian cells. In turn, nuclear import capacity and nuclear size scale as a function of ELYS expression. In addition to identifying a novel modulator of nuclear size, this is, to our knowledge, the first demonstration that NPC density and nuclear import capacity impact nuclear size in mammalian cells.

## RESULTS

### A high-throughput imaging-based siRNA screen identifies ELYS as a nuclear size effector

We carried out a high-throughput imaging RNAi screen in a premalignant breast epithelial cell line (MCF-10AT1k.cl2) to identify factors that affect nuclear size (Fig. 1A), with an emphasis on factors whose loss results in smaller nuclei. Briefly, cells were transfected in 384-well format with siRNA oligo libraries targeting a total of 867 genes (three unique siRNA oligo sequences per gene) implicated in NE function, chromatin structure, and epigenetic mechanisms (for details see Methods). The screen was performed in two biological replicates. As a positive control, lamin B1 (LMNB1) was knocked down to decrease nuclear size [36], and a non-targeting siRNA was used as a negative control on each plate (Fig. 1B and 2A). 48 h after siRNA oligo transfection, cells were fixed, stained for DNA and nuclear lamins, and imaged using high-throughput confocal microscopy (see Methods). Automated high content image analysis generated measurements of nuclear cross-sectional area, a reliable proxy for detecting changes in nuclear volume (Fig. S1A) [30,32,42,54]. Statistical analysis of the image analysis output was performed, and genes for which silencing with at least 2 out of the 3 siRNA oligos led to a z-score of <-1.5 (i.e. decreased nuclear size) were classified as putative nuclear size effectors (see Methods).

**Figure 1:**
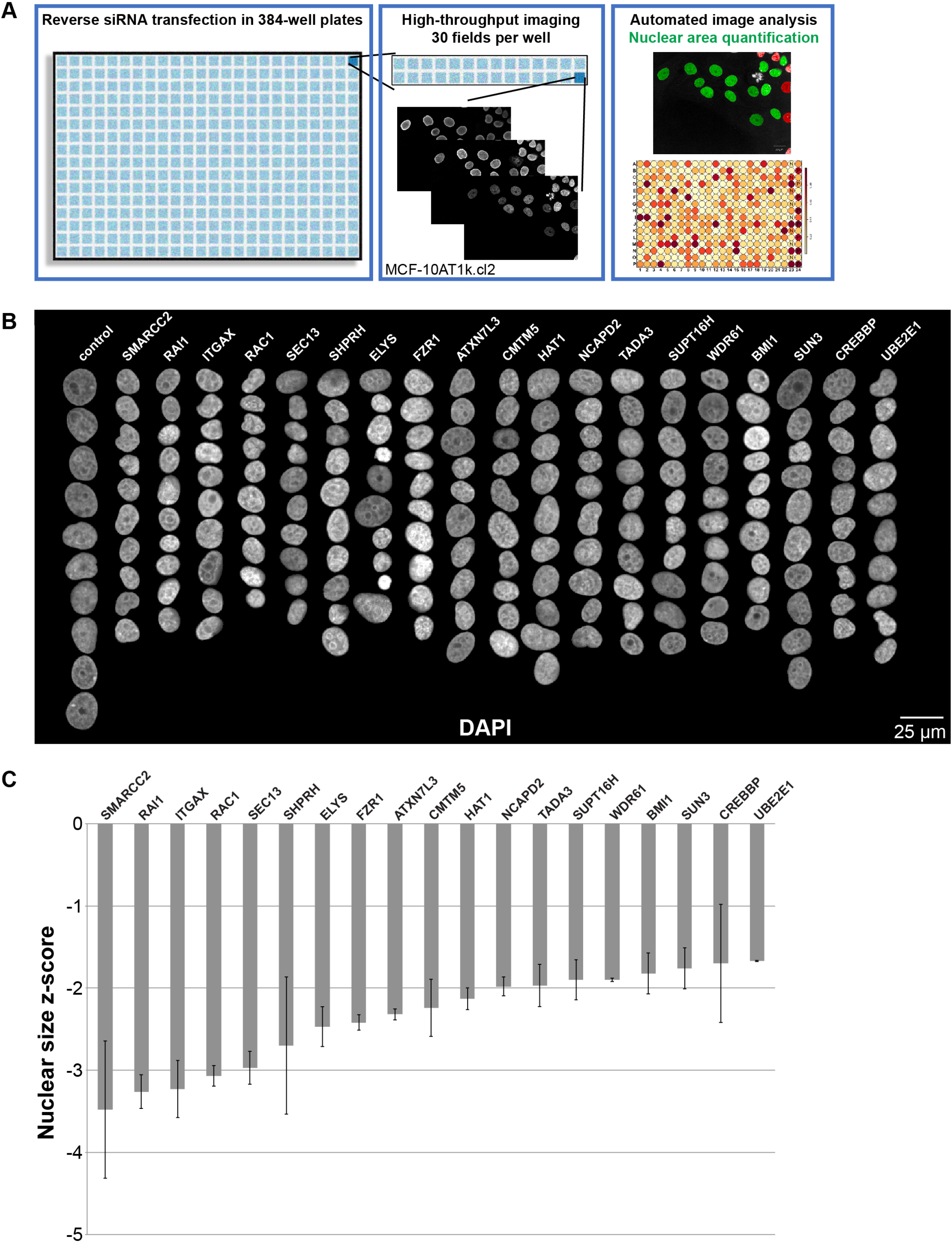
An imaging-based siRNA screen for gene knockdowns that reduce nuclear size. (**A**) The screen approach is depicted, and details are available in the Methods. Throughout the study, a premalignant breast cancer cell line was used (MCF-10AT1k.cl2), unless otherwise indicated. Images and data shown in this figure were obtained directly from the screen. (**B**) Montages of representative DAPI-stained nuclei are shown for gene knockdowns that led to reduced nuclear size. (**C**) Maximum nuclear size z-scores are plotted for gene knockdowns that led to reduced nuclear size. Data are based on three different siRNA oligo sequences for each gene and two biological replicates. Error bars represent SEM.

**Figure 2:**
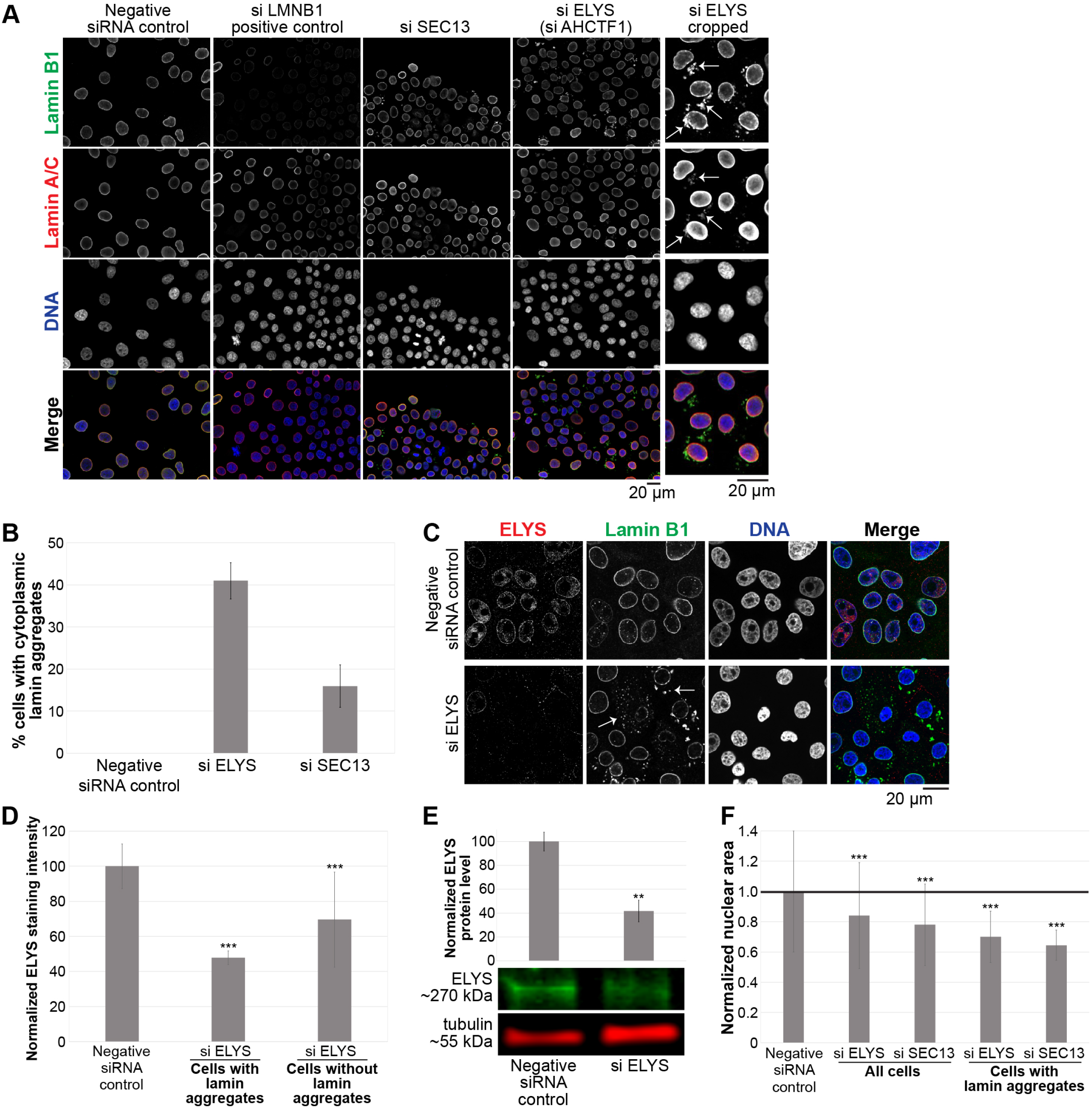
ELYS and SEC13 knockdown lead to smaller nuclei and the formation of cytoplasmic lamin aggregates. (**A**) Representative images from the screen are shown. To the far right, ELYS knockdown nuclei were cropped to highlight cytoplasmic lamin aggregates indicated with arrows. (**B**) Cells with and without cytoplasmic lamin B1 aggregates were counted and the percentage of cells with aggregates was calculated. For each experiment, 698-1213 cells were examined per condition. Data are shown for two siRNA sequences for each gene and two biological replicates. (C-D) Cells were transfected with control or ELYS siRNA and stained with ELYS and lamin B1 antibodies as indicated. Representative images are shown. Some examples of cytoplasmic lamin aggregates are indicated with arrows. For each experiment, nuclear ELYS staining intensity was quantified for 90-143 nuclei per condition and normalized to the negative control. Data from two biological replicates are shown. (**E**) Cell lysates from control and ELYS siRNA transfected cells were analyzed by western blot and probed for ELYS and tubulin. One representative western blot is shown. ELYS band intensity was normalized to tubulin. Quantification from two biological replicates is shown. (**F**) Nuclear cross-sectional areas were quantified for 70-762 nuclei per condition, averaged, and normalized to the negative control. One representative experiment of six is shown. ***, p<0.005; **, p<0.01; error bars represent SD.

Out of 867 genes screened, knockdown of 19 resulted in decreased nuclear size with median z-scores <-1.5 (Table S1–S2, Fig. 1B-C). The Nup ELYS was identified as a strong hit with a median z-score of −2.0, in the top ~30% of hits. ELYS is a component of the Nup107-160 complex that has important roles in NPC assembly [55–60]. ELYS was of particular interest given its role in nuclear pore formation, hence implicating nuclear transport in nuclear size regulation, and also because its siRNA knockdown not only induced smaller nuclei but also resulted in formation of cytoplasmic lamin puncta, containing both A- and B-type lamins, in up to 40% of cells (Fig. 2A-B; see below). The frequency of cells with cytoplasmic lamin aggregates correlated with higher ELYS knockdown (Fig. 2C-D). Near complete knockdown of ELYS in HeLa cells was previously reported to cause cytokinesis defects [49], however we did not observe obvious cell cycle effects (Fig. S1C), possibly because overall ELYS proteins levels were typically only reduced by ~60% (Fig. 2E) and/or because we are studying a breast epithelial cell line rather than a cancer cell line. It is likely that partial ELYS knockdown was important in allowing us to observe and characterize the nuclear size phenotype. The screen also identified another component of the ELYS containing Nup107-160 complex, SEC13 [61,62], the top hit with a median z-score of −2.7. Similar to ELYS, knockdown of SEC13 induced smaller nuclei and cytoplasmic lamin puncta (Table S1–S2, Fig. 1B-C and 2A-B). While not all ELYS and SEC13 knockdown cells exhibited lamin puncta, those with puncta had smaller nuclei compared to knockdown cells without puncta (Fig. 2F). This suggests that accumulation of lamins in the cytoplasm is associated with smaller nuclei.

The observed effect of ELYS and SEC13 is specific and not a general property of Nups since, out of 33 Nups tested in the screen, knockdown of only these two Nups decreased nuclear size. While ELYS and SEC13 knockdown might be expected to have pleiotropic effects, there was no pronounced change in the cell cycle profile (Fig. S1C), suggesting that observed nuclear size reductions were not indirectly due to altered cell proliferation or cell cycle progression. Furthermore, in three different cell lines (human dermal fibroblasts, MCF-10A normal breast epithelial cells, and MCF-10AT1k.cl2 premalignant breast epithelial cells), ELYS and SEC13 knockdown led to smaller nuclei and, to varying degrees, cytoplasmic lamin puncta (Fig. S2). Because SEC13 plays dual roles in NPC assembly and protein trafficking, we focused our subsequent analysis on ELYS.

### Cytoplasmic lamin puncta contain multiple lamin types and are dynamic

ELYS knockdown cells stained for various lamin types revealed that cytoplasmic lamin puncta contain lamin B1, B2, and A/C (Fig. 3A-C). Nuclear levels of lamins A/C and B1 appeared unaltered, whereas lamin B2 nuclear levels were reduced by about 30% (Fig. 3A-C; p < 0.005). To confirm that lamin puncta were not an artifact of fixation, we transfected cells with an mCherry-lamin B2 expression construct and performed live cell imaging. Cytoplasmic lamin puncta were apparent in ELYS knockdown cells but not in control cells (Fig. 3D). In addition, time-lapse imaging showed that cytoplasmic lamin aggregates exhibited dynamic movements, often appeared immediately after mitosis regardless of whether or not the mother cell contained lamin puncta, and coalesce or dissolve during interphase (Movies 1-5). In some extreme cases, cells contained abundant cytoplasmic lamin foci but no NE-localized mCherry-lamin B2 (Fig. 3D, Movies 2 and 6).

**Figure 3:**
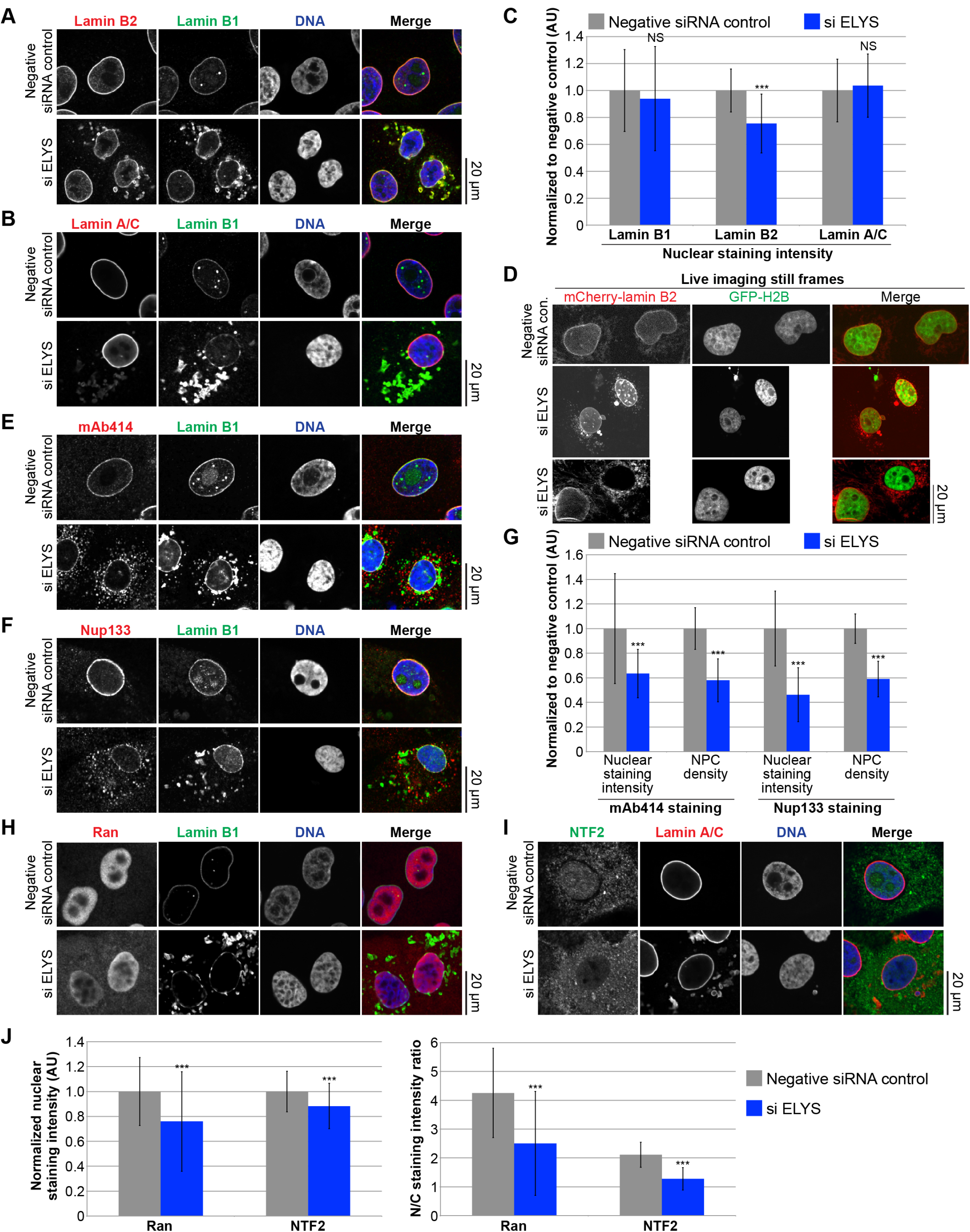
ELYS knockdown reduces NPC density and nuclear levels of lamin B2, Ran, and NTF2. MCF-10AT1k.cl2 cells were transfected with control or ELYS siRNA and stained with antibodies against: (**A**) lamin B2 and B1, (**B**) lamin A/C and B1, (**E**) FG-Nups (mAb414) and lamin B1, (**F**) Nup133 and lamin B1, (**H**) Ran and lamin B1, (**I**) NTF2 and lamin A. Representative images are shown. (**C**) For each experiment, nuclear lamin B1, B2, and A/C staining intensities were quantified for 44-159 nuclei per condition (96 nuclei on average) and normalized to the negative control. Data from one representative experiment of three are shown. (**D**) MCF-10AT1k.cl2 cells were cotransfected with control or ELYS siRNA and with plasmids expressing mCherry-lamin B2 and H2B-GFP. Representative images of live cells are shown. Also see Movies 1-6. (**G**) For each experiment, nuclear mAb414 and Nup133 staining intensities were quantified for 31-83 nuclei per condition (60 nuclei on average) and normalized to the negative control. To measure NPC densities, confocal NE surface images were acquired for mAb414 and Nup133 stained nuclei. NPC numbers were counted per unit area for 18-50 nuclei per condition (30 nuclei on average) and normalized to the negative control. Data from one representative experiment of two are shown. (**J**) Nuclear Ran and NTF2 staining intensities were quantified and normalized to the negative control. To measure the N/C staining intensity ratio of a given cell, the average fluorescence intensity of a nuclear region was divided by the average fluorescence intensity of a cytoplasmic region. For each experiment, 54-216 nuclei were quantified per condition (112 nuclei on average). Data from one representative experiment of two are shown. ***, p<0.005; NS, not significant; error bars represent SD.

Because lamins are known to be regulated by phosphorylation [32,63–65], we asked whether the appearance of cytoplasmic lamin aggregates might depend on the lamin phosphorylation state. To test this hypothesis, we knocked down two putative lamin protein phosphatases, PPP1CA and PPP2R4 [66]. While over 40% of ELYS knockdown cells exhibited cytoplasmic lamin puncta, less than 10% of cells cotransfected with siRNA against ELYS and PPP1CA showed puncta (Fig. S3A-B). Knockdown of PPP2R4 had no impact on the appearance of lamin puncta (data not shown). Dissolving cytoplasmic lamin aggregates through PPP1CA knockdown did not result in a significant increase in nuclear size in ELYS knockdown cells (Fig. S3C). Thus, cytoplasmic lamin aggregates formed in ELYS knockdown cells are not sufficient to cause reduced nuclear size.

### ELYS levels control NPC density and nuclear localization of Ran and NTF2

To begin to address the mechanism by which ELYS influences nuclear size, we stained cells for FG-Nups and Nup133, another component of the Nup107-160 complex, to assess nuclear pore number and integrity. Nuclear Nup staining intensity and NPC density were both reduced in ELYS knockdown cells (Fig. 3E-G), correlating with reduced nuclear localization of Ran and NTF2 (Fig. 3H-J). Because NTF2 associates with the NPC and mediates Ran import [67–73], reduced levels of nuclear Ran and NTF2 likely reflect reduced NPC numbers, consistent with our observation of reduced nuclear lamin B2 (Fig. 3A-C). We also noted cytoplasmic staining for FG-Nups and Nup133 in ELYS knockdown cells that was not evident in control cells and did not appear to colocalize with lamin puncta (Fig. 3E-F). Co-transfections with ELYS siRNA and a plasmid expressing an siRNA-resistant form of GFP-ELYS rescued the effects of ELYS knockdown alone, resulting in increased nuclear size, nuclear lamin B2 staining, and NPC density as well as a dramatic reduction in the percentage of cells with cytoplasmic lamin aggregates (Fig. 4A-D, G-H). These data indicate that the phenotypes observed for ELYS knockdown cells are not due to off-target effects.

**Figure 4:**
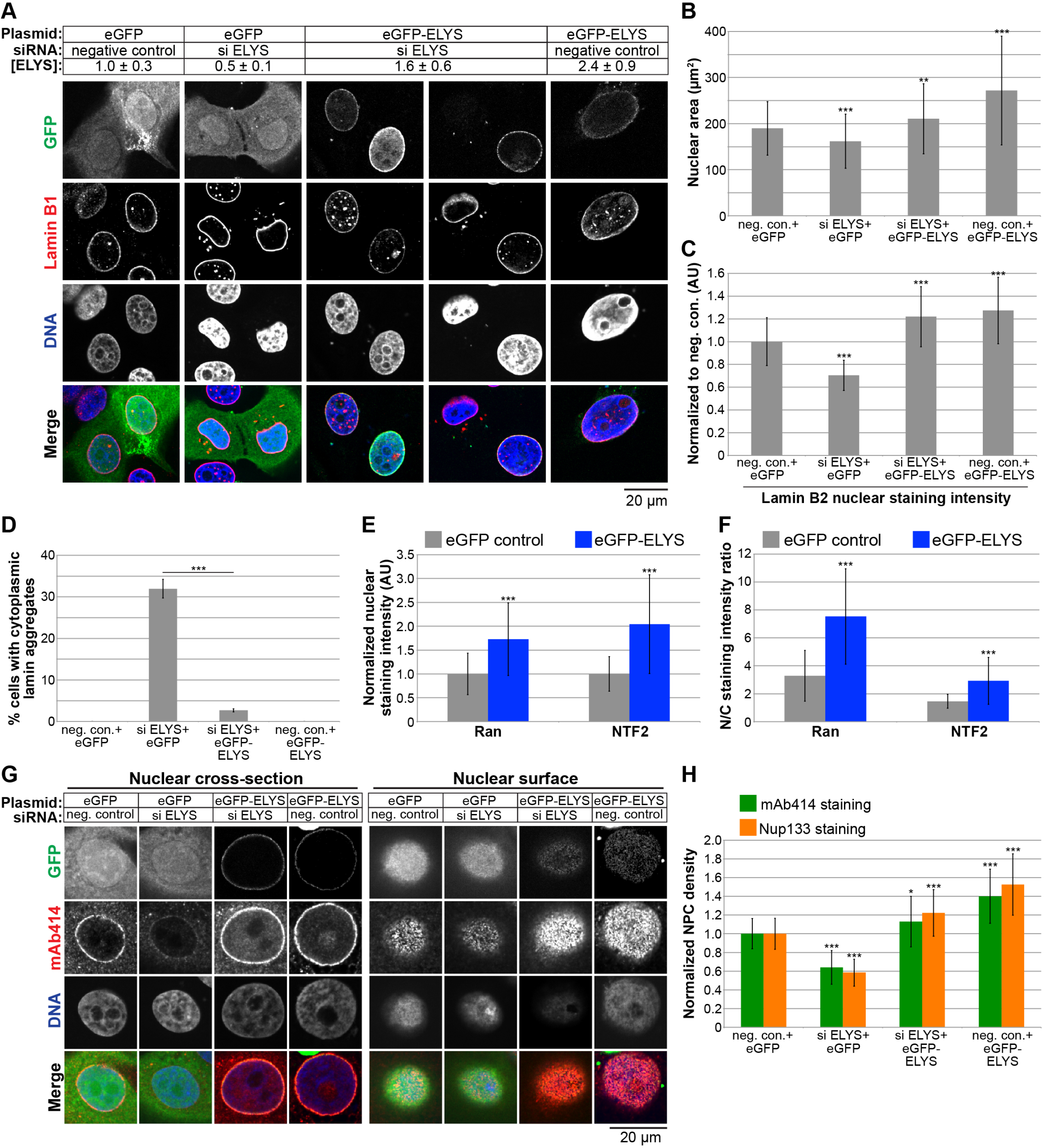
ELYS overexpression increases nuclear size, NPC density, and nuclear levels of lamin B2, Ran, and NTF2. (**A**) MCF-10AT1k.cl2 cells were co-transfected with control or ELYS siRNA and plasmids expressing eGFP alone or siRNA-resistant eGFP-ELYS. Western blots were performed and quantified as described in Figure 2E to quantify ELYS concentrations, normalized to the eGFP/negative siRNA control and indicated in the row labeled [ELYS] (average ± SD). Quantification was performed on two biological replicates. Cells were stained with an antibody against lamin B1. Representative images are shown. (**B**) For each experiment, nuclear cross-sectional areas were quantified for 68-298 nuclei per condition (162 nuclei on average) and averaged. Data from one representative experiment of three are shown. (**C**) For each experiment, nuclear lamin B2 staining intensities were quantified for 29-81 nuclei per condition (60 nuclei on average) and normalized to the eGFP/negative siRNA control. Data from one representative experiment of two are shown. (**D**) Cells with and without cytoplasmic lamin B2 aggregates were counted and the percentage of cells with aggregates was calculated. For each experiment, 58-299 cells were examined per condition. Data from one representative experiment of three are shown. (**E-F**) Cells were stained with antibodies against Ran and NTF2. For each experiment, 62-130 nuclei were quantified per condition (92 nuclei on average). Data from one representative experiment of two are shown. (**E**) Nuclear Ran and NTF2 staining intensities were quantified and normalized to the eGFP control. (**F**) To measure the N/C staining intensity ratio of a given cell, the average fluorescence intensity of a nuclear region was divided by the average fluorescence intensity of a cytoplasmic region. (**G**) Transfections were performed as in (A) and cells were stained with an antibody against FG-Nups (mAb414) and Nup133. Representative images for mAb414 staining are shown. Confocal imaging was performed through nuclear cross-sections as well as on the surface of nuclei. (**H**) To measure NPC densities, confocal NE surface images were acquired for mAb414 and Nup133 stained nuclei. NPC numbers were counted per unit area for 25-116 nuclei per condition (53 nuclei on average) and normalized to the negative control. Data from one representative experiment of two are shown. ***, p<0.005; **, p<0.01; *, p<0.05; error bars represent SD.

We also examined the effect of ELYS overexpression by transfecting cells with GFP-ELYS (Fig. 4A). ~2.4-fold ELYS overexpression led to a greater than 40% increase in nuclear cross-sectional area with no apparent lamin aggregates (Fig. 4A-D). Furthermore, NPC density was increased in ELYS overexpressing cells, along with a concomitant increase in nuclear localization of Ran, NTF2, and lamin B2 (Fig. 4C, E-H). Taken together, these data show that modulating ELYS levels affects NPC numbers and the localization of nuclear transport factors and lamins, suggesting that changes in nuclear import capacity underlie observed changes in nuclear size. We believe this is the first demonstration that NPC densities can impact nuclear size.

### Nuclear import capacity tunes nuclear size

Because ELYS knockdown results in reduced NPC density that might abate nuclear import capacity, we tested if increasing bulk import in ELYS knockdown cells would rescue the observed nuclear size defect. Indeed, overexpression of importin α in ELYS knockdown cells resulted in nuclear sizes comparable to control cells and a reduction in the percentage of cells with cytoplasmic lamin puncta (Fig. 5A-C). These data indicate that small nuclear size and the formation of lamin aggregates in ELYS knockdown cells are due to limited nuclear import capacity. Consistent with this notion, importin α overexpression increased nuclear levels of both lamin B2 and GFP-3x SV40 NLS, a reporter of importin α/β-mediated nuclear import (Fig. 5D). We also observed that importin α overexpression alone resulted in a 40% increase in nuclear cross-sectional area compared to control cells (Fig. 5A-B), similar to what we observed in HeLa, U2OS, and MRC5 cells (data not shown).

**Figure 5:**
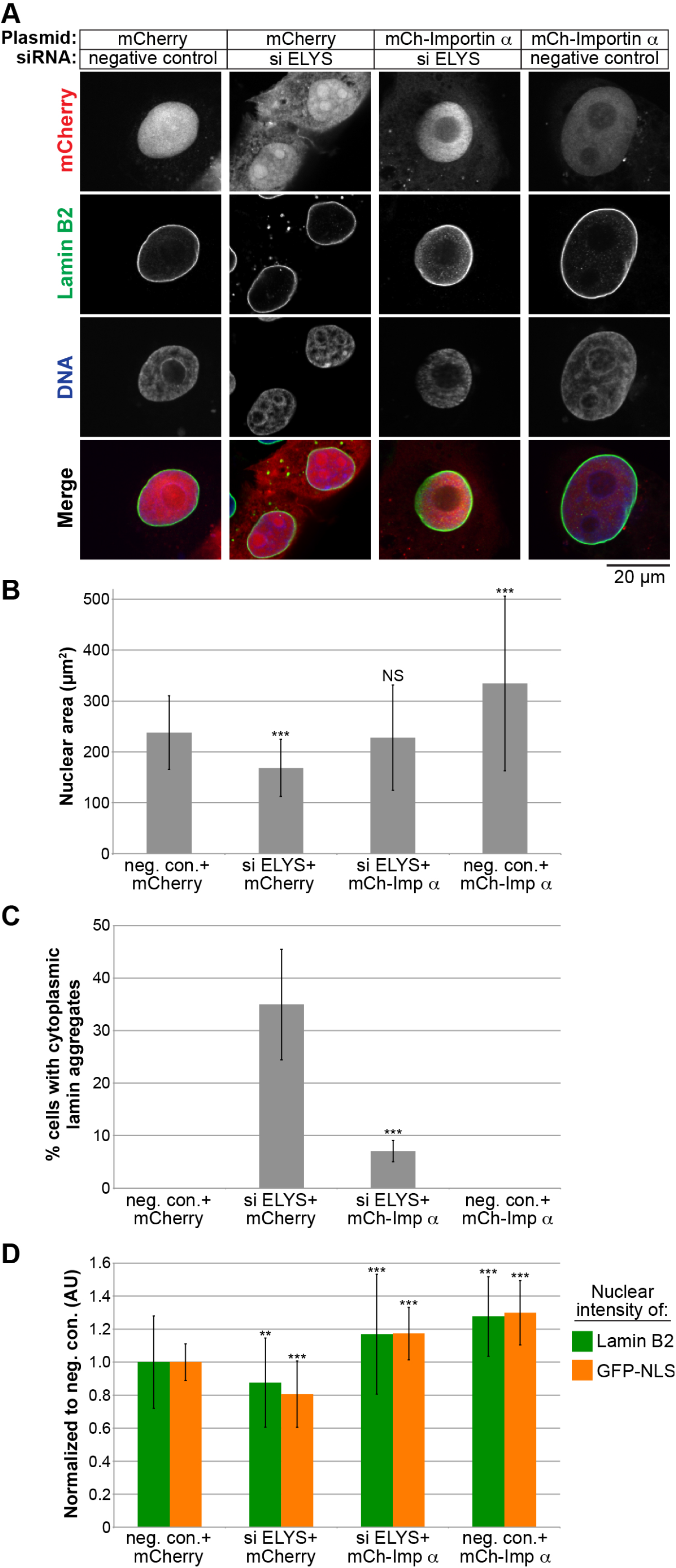
Increasing nuclear import by overexpressing importin α rescues nuclear size and lamin aggregation phenotypes in ELYS knockdown cells. (**A**) MCF-10AT1k.cl2 cells were co-transfected with control or ELYS siRNA and plasmids expressing mCherry alone or mCherry-importin a. Cells were stained with an antibody against lamin B2. Representative images are shown. (**B**) For each experiment, nuclear cross-sectional areas were quantified for 30-172 nuclei per condition (84 nuclei on average) and averaged. Data from one representative experiment of two are shown. (**C**) Cells with and without cytoplasmic lamin B2 aggregates were counted and the percentage of cells with aggregates was calculated. For each experiment, 74-218 cells were examined per condition. Data from one representative experiment of two are shown. (**D**) For each experiment, nuclear lamin B2 staining intensities were quantified for 34-187 nuclei per condition (85 nuclei on average) and normalized to the mCherry/negative siRNA control. To measure bulk import capacity for importin α/β cargos, cells were co-transfected as in (**A**) along with a plasmid expressing GFP-3x SV40 NLS. For each experiment, nuclear GFP-NLS intensities were quantified for 22188 nuclei per condition (73 nuclei on average) and normalized to the mCherry/negative siRNA control. Data from one representative experiment of two are shown. ***, p<0.005; **, p<0.01; NS, not significant; error bars represent SD.

To test more generally the impact of nuclear import capacity on nuclear size we inhibited nuclear import in two ways: 1) by expressing RanQ69L that is constitutively GTP bound and acts as a dominant negative inhibitor of importin β-mediated nuclear import [74,75], and 2) by treating cells with importazole [76], a small molecule inhibitor of the interaction between importin β and RanGTP (Fig. 6A-B). Both experimental approaches resulted in reduced bulk import capacity for importin α/β cargos, as expected, with concomitant reductions in nuclear size and the appearance of cytoplasmic lamin aggregates in a subset of cells (Fig. 6C-E). Taken together, these data show that limiting import capacity either by reducing NPC density or targeting nuclear transport factors results in reduced nuclear size and the cytoplasmic accumulation of nuclear lamin proteins that have the propensity to aggregate.

**Figure 6:**
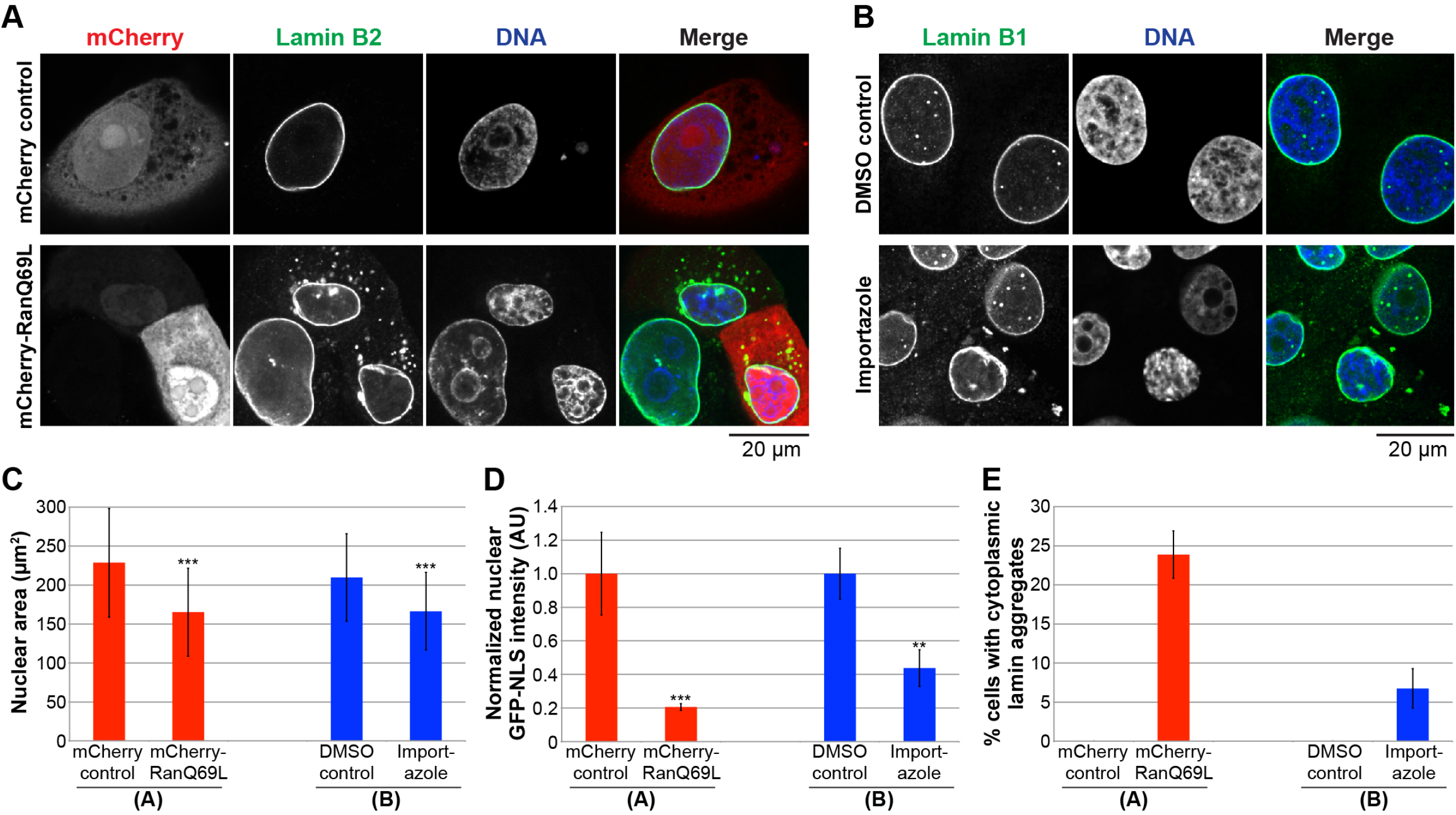
Inhibiting nuclear import leads to smaller nuclei and cytoplasmic lamin aggregates. (**A**) MCF-10AT1k.cl2 cells were transfected with plasmids expressing mCherry alone or mCherry-RanQ69L and stained with an antibody against lamin B2. Representative images are shown. (**B**) MCF-10AT1k.cl2 cells were treated with 20 μM importazole or an equivalent volume of DMSO as a control for 24 hours and stained with an antibody against lamin B1. Representative images are shown. (**C**) For each experiment, nuclear cross-sectional areas were quantified for 32-271 nuclei per condition (123 nuclei on average) and averaged. Data from one representative experiment of three are shown. (**D**) To measure bulk import capacity for importin α/β cargos, experiments were performed as in (A-B) except that cells were also transfected with a plasmid expressing GFP-3x SV40 NLS. For each experiment, nuclear GFP-NLS intensities were quantified for 11-75 nuclei per condition (35 nuclei on average) and normalized to the appropriate control. Data from one representative experiment of two are shown. (**E**) Cells with and without cytoplasmic lamin B aggregates were counted and the percentage of cells with aggregates was calculated. For each experiment, 31-130 cells were examined per condition. Data from one representative experiment of three are shown. ***, p<0.005; **, p<0.01; error bars represent SD.

Given that reducing nuclear import decreased nuclear size, we predicted that inhibiting nuclear export might lead to an increase in nuclear size. We first knocked down expression of the nuclear export factor XPO1 [77–79], which increased nuclear size and nuclear levels of lamins B1 and A/C (Figs. S1A-B, S2, and S4A-C), without grossly perturbing cell cycle progression (Fig. S1C). Knockdown of lamin B1 and A in combination with XPO1 slightly reduced nuclear size, but not to the size of nuclei in control cells, indicating that increased lamin import in XPO1 knockdown cells cannot fully account for the increased nuclear size (Fig. S4C). Indeed, staining for total protein showed increased accumulation of nuclear proteins in XPO1-depleted cells (Fig. S4D-E). Consistent with these data, inhibition of XPO1 with the small molecule leptomycin B also increased nuclear size and total nuclear protein levels (Fig. S4D-E). These data suggest that blocking nuclear export leads to increased accumulation of proteins in the nucleus, potentially driving increased nuclear size. Furthermore, XPO1 knockdown increased nuclear size to varying degrees in four different cells lines (Fig. S2). Taken together, these results are consistent with a model in which modulating protein transport across the NPC represents a key mechanism of nuclear size control. We believe this is the first demonstration in mammalian cells that nuclear transport capacity can tune nuclear size.

## DISCUSSION

We have used an imaging-based high-throughput RNAi screen for nuclear size effectors to identify ELYS as a novel modulator of mammalian nuclear size. Out of 867 screened genes, knockdown of only 19 decreased nuclear size. Of those, two Nups were identified (ELYS and SEC13), whereas knockdown of 31 other Nups did not decrease nuclear size, demonstrating specificity of the hits. Toward mechanism, we showed that decreasing ELYS expression levels resulted in a concomitant decrease in NPC density, importin α/β-mediated nuclear import, and nuclear size, while ELYS overexpression increased NPC density, nuclear import, and nuclear size. These data support a scenario whereby NPC number and nuclear import capacity scale nuclear size. Consistent with this model, we find that modulating nuclear import capacity alters nuclear size. One particularly striking observation was that importin α overexpression was sufficient to overcome the NPC density defect in ELYS knockdown cells, highlighting how import capacity is dictated by a delicate balance between NPC numbers and transport factor levels. Furthermore, inhibiting nuclear export in favor of import led to larger nuclei, suggesting that there is competition between import and export through limited numbers of NPCs. While the identities of the imported cargos required for nuclear growth are still unknown, nuclear lamins have previously been implicated in this process [30,34–36,80,81]. Our data also support a role for lamins, as their nuclear levels were decreased upon ELYS knockdown and increased upon XPO1 knockdown, and lamin knockdown mitigated some of the nuclear growth observed in XPO1 depleted cells. The observation that total nuclear protein levels were increased when nuclear export was inhibited is consistent with findings in fission yeast [27,28].

Our study is, to our knowledge, the first demonstration that NPC numbers can modulate nuclear size and that nuclear transport can tune nuclear size in mammalian cells. Experiments using *Xenopus* egg extract and early embryos suggested that NPC numbers are not limiting for nuclear import or nuclear size [30,82,83], likely because nuclear import capacity is extremely high in these systems. In HeLa cells, cyclin-dependent kinase inhibition blocked interphase NPC assembly leading to a reduction in NPC density, yet nuclear growth was unaffected [84,85], and ELYS depletion did not significantly affect nuclear lamin localization or import capacity [86]. These results might be explained by an upregulation of nuclear import that is frequently observed in cancer cells [87–90]. These studies highlight the importance of cell type and disease state when considering mechanisms of nuclear size control. Notably, NPC densities may dominate nuclear size control in normal somatic cells while transport factor levels and activities may play a more important role in early development and cancer. Consistent with this idea, ELYS, SEC13, and XPO1 knockdown had variable effects on nuclear size in different cell lines (Fig. S2). In particular, ELYS and SEC13 knockdown significantly reduced nuclear size in three roughly normal cell lines but minimally affected nuclear size in MCF7 breast cancer cells (Fig. S2). In summary, we propose that nuclear size is determined by import capacity, which in turn is controlled by NPC density and levels of nuclear transport factors.

One striking and unexpected phenotype of ELYS knockdown was the appearance of lamin puncta in the cytoplasm. As lamin foci were more prevalent in cells with lower ELYS protein levels, we suspect that reduced nuclear import of lamins leads to cytoplasmic lamin accumulation and aggregation. This does not appear to be a phenotype specific to ELYS knockdown because inhibiting nuclear import by RanQ69L expression or importazole treatment also resulted in a subset of cells with lamin puncta. These data suggest that sufficiently high lamin import kinetics are necessary to avoid cytoplasmic aggregation. Time-lapse imaging revealed that lamin puncta are dynamic and move, potentially impacting cytoplasmic activities. While it is unknown if these cytoplasmic lamin aggregates affect cell function, they may be relevant to physiological conditions or disease states where nuclear import is compromised. For example, defective nuclear transport is linked to amyotrophic lateral sclerosis, frontotemporal dementia, and Huntington’s disease [91–95]. Furthermore, cytoplasmic lamin puncta may contribute to the pathology of diseases associated with lamin mislocalization, as is observed in some laminopathies [96–101]. For instance, mislocalization of lamin A to the cytoplasm results in the formation of lamin aggregates that sequester emerin away from the nucleus [102]. In addition, cellular stress induces the aggregation of nucleocytoplasmic transport factors into cytoplasmic stress granules that are implicated in the pathogenesis of several neurodegenerative diseases [103]. Going forward, it will be interesting to investigate what other nuclear proteins accumulate in the cytoplasm when import is compromised and whether these proteins engage in inappropriate and potentially pathogenic interactions.

In conclusion, using an siRNA screening approach, we have uncovered a novel mammalian nuclear size effector. Changes in nuclear size are commonly used in cancer diagnosis and prognosis, yet it is unknown whether altered nuclear size contributes to, or is simply a consequence of, the pathology. Levels of nuclear transport factors are frequently altered in cancer and have begun to be targeted for cancer treatments [10,87,88,90,104–106]. Our results suggest that it may also be important to examine changes in NPC densities in cancer.

## MATERIALS AND METHODS

### Cell culture and small molecule treatments

The MCF-10AT1 k.cl2 and MCF-10A cell lines were obtained from the Barbara Ann Karmanos Cancer Institute [107–109]. MCF-10AT1k.cl2 cells were passaged every ~3 days and were cultured at 37°C and 5% CO_2_ in DMEM/F12 (1:1), 1.05 mM CaCl_2_, 4.9% horse serum, 10 mM HEPES, 10 μg/ml insulin, 20 ng/ml EGF, and 0.5 μg/ml hydrocortisone. MCF-10A cells were passaged every ~5 days and were cultured at 37°C and 5% CO2 in DMEM/F12 (1:1), 1.05 mM CaCl_2_, 4.9% horse serum, 10 mM HEPES, 10 μg/ml insulin, 20 ng/ml EGF, 0.5 μg/ml hydrocortisone, and 0.1 μg/ml cholera enterotoxin. We used previously described hTERT immortalized dermal fibroblast cells [110], maintained in MEM supplemented with 15% fetal bovine serum, 2 mM L-glutamine, 1 mM sodium pyruvate, 100 U/mL penicillin, and 100 μg/mL streptomycin. The MCF7 cell line was obtained from ATCC and cultured in MEM supplemented with 100 μg/ml insulin, 10% fetal bovine serum, 100 U/mL penicillin, and 100 μg/mL streptomycin. Cells were cultured at 37°C and 5% CO_2_. Importazole (MedChemExpress # HY-101091) was dissolved in DMSO and cells were treated with a final concentration of 20 μM for 24 hours. Leptomycin B (Santa Cruz Biotechnology # sc-358688) was dissolved in ethanol and cells were treated with a final concentration of 20 ng/ml for 24 hours.

### Plasmids and siRNA sequences

The following plasmids were obtained commercially: pEGFP-C2 (Clontech #6083-1), pcDNA-EGFP-ELYS-polyA (Addgene #59746) was a gift from Yi Zhang [111], pCMV/myc/nuc/GFP which consists of GFP fused to 3x SV40 nuclear localization signals whose nuclear import is mediated by importin α/β (Invitrogen V821-20), pmCherry-C1-RanQ69L (Addgene #30309) was a gift from Jay Brenman [112], H2B-GFP (Addgene #11680) was a gift from Geoff Wahl [113]. Plasmid pmCherry-C2 is a derivative of pEGFP-C2 and was a gift from Anne Schlaitz. Plasmid pmCherry-C2-lamin B2 (pDL24) was described previously [36]. Plasmid pmCherry-C2-importin alpha was constructed by cloning hSRP1a into pmCherry-C2 at EcoRI and KpnI (pDL21). The following siRNAs were ordered from Life Technologies:

ELYS (siRNA ID # s24727): sense 5’ GCGAUUGUCUGCUUACAGAtt 3’, antisense 5’ UCUGUAAGCAGACAAUCGCtc 3’ (note that this siRNA does not target ELYS in pcDNA-EGFP-ELYS-polyA)
SEC13 (siRNA ID # s12662): sense 5’ CAAUUACAUCAAGAGGUUUtt 3’, antisense 5’ AAACCUCUUGAUGUAAUUGgg 3’
PPP1CA (siRNA ID # s10930): sense 5’ CAUCUAUGGUUUCUACGAUtt 3’, antisense 5’ AUCGUAGAAACCAUAGAUGcg 3’
PPP2R4 (siRNA ID # s10978): sense 5’ GGAUUCAUCCUUACCCUCAtt 3’, antisense 5’ UGAGGGUAAGGAUGAAUCCga 3’
XPO1 (siRNA ID # s14939): sense 5’ CCAAUAUUCGACUUGCGUAtt 3’, antisense 5’ UACGCAAGUCGAAUAUUGGta 3’
LMNA (siRNA ID # s8222): sense 5’ GAAGGAGGGUGACCUGAUAtt 3’, antisense 5’ UAUCAGGUCACCCUCCUUCtt 3’
LMNB1 (siRNA ID # s8225): sense 5’ GAAUCGUUGUCAGAGCCUUUU 3’, antisense 5’ AAGGCUCUGACAACGAUUCUC 3’
Negative control No. 1 si RNA (catalog number 4390843)

### Transfections

For siRNA transfection, reverse transfection was performed in 6- or 24-well plates using Lipofectamine RNAiMAX (ThermoFisher Scientific) with a final siRNA concentration of 20 nM according to the manufacturer’s protocol. For plasmid transfection, reverse transfection was performed in 24-well plates using Lipofectamine 3000 (ThermoFisher Scientific) with 500 ng of plasmid per well according to the manufacturer’s protocol. For co-transfection of siRNA and plasmid, reverse transfection was performed in 24-well plates using Lipofectamine 3000 (ThermoFisher Scientific) with 500 ng of plasmid per well and a final siRNA concentration of 30 nM according to the manufacturer’s protocol.

### High-throughput siRNA oligo transfections

MCF-10AT1k.cl2 cells were reverse transfected with the oligo siRNA libraries (siRNA Silencer Select, ThermoFisher), positive control (LMNB1, ThermoFisher, s8225), and negative non-targeting control (ThermoFisher cat#4390847) in 384-well imaging plates (CellCarrier-384 PerkinElmer, 6057300). The RNAi libraries used were a custom library targeting gene products predicted to localize to the nuclear envelope (346 genes, ThermoFisher, lot# AMO20JUZ), and an off the shelf library targeting proteins involved in epigenetic and chromatin regulation (521 genes, ThermoFisher, cat# A30085, lot# AMO20K2X). Each gene was targeted with 3 individual siRNAs, for a total of 2601 siRNAs in the screen. Assay ready plates were prepared by first spotting 2 μl of a 400 nM solution of siRNA oligos at the bottom of a dry 384-well plate using the MDT module of a Janus automated liquid handler (PerkinElmer), and then air-drying under a laminar sterile air-flow for 2 hrs. For cell cycle experiments (Fig. S1C) and experiments with different cell lines (Fig. S2), 150 nl of an siRNA oligo solution at 5 μM concentration were spotted at the bottom of imaging plates using an Echo525 instrument (Labcyte). In these cases, the negative control No. 1 si RNA (Life Technologies, catalog number 4390843) non-targeting siRNA control was used. The assay ready plates were then sealed and stored at −80°C until the day of the transfection, when they were thawed, equilibrated at room temperature, and centrifuged at 1000rpm for 20 min. 20 μl of a solution containing 0.035 μl of RNAiMax transfection reagent (ThermoFisher, cat#13778) were dispensed in each well of the assay ready plates using a Multidrop Combi Reagent Dispenser (ThermoFisher). The siRNA oligo/RNAiMax solution was then incubated at room temperature for 20 min. 20 μl of a cell suspension containing 1200 cells in DMEM/F12 (1:1), 1.05 mM CaCl_2_, 9.8% horse serum, 10 mM HEPES, 10 μg/ml insulin, 20 ng/ml EGF, and 0.5 μg/ml hydrocortisone were then dispensed in each well of the plate using a Multidrop Combi Reagent Dispenser. The final concentration of the siRNA oligos in each well was 20 nM. Cells were then incubated for 48 hrs at 37°C. The screen was performed in two biological replicates on different days. Cell cycle experiments (Fig. S1C) and experiments with different cell lines (Fig. S2) were performed in 3 technical replicates over 3 independent biological replicates, for a total of 9 wells per experimental condition.

### High-throughput immunofluorescence

siRNA oligo transfected cells were fixed in 4% PFA in PBS for 20 min at room temperature, washed 3 × 5 min in PBS, permeabilized with 0.5% Triton-100x in PBS for 15 min, washed 3 × 5 minutes in PBS, and blocked in PBS with 0.05% Tween 20 (PBST) and 5% BSA. To visualize the periphery of the cell nucleus, cells were immunostained with primary antibodies against lamin A/C (Santa Cruz, sc-376248, mouse, 1:1000) and lamin B1 (Santa Cruz, sc-6217, goat, 1:500) in PBST with 1% BSA for 4 hours at room temperature or overnight at 4°C. Cells were washed 3 × 5 minutes with PBST and incubated for one hour at room temperature with secondary antibodies diluted in 1% BSA in PBST containing DAPI (5 ng/μl). Secondary antibodies were 1:500 dilutions of Alexa Fluor 488 anti-goat IgG (Molecular Probes, A-11055) and Alexa Fluor 568 anti-mouse IgG (Molecular Probes, A-11031). Plates were then washed 3 × 5 minutes in PBST, sealed, and stored at 4°C until imaging. All the automated liquid handling steps necessary for high-throughput immunofluorescence staining of the 384-well plates were performed using a Biotek EL406 plate washer.

### High-throughput image acquisition

Image acquisition was performed using an Opera QEHS (PerkinElmer) high-throughput dual spinning disk confocal microscope. Images were acquired using the 40X water immersion lens (N.A. 0.9) and two CCD cameras (1.3 MPixels) with pixel binning of 2 (Pixel size: 323 nm). For the DAPI channel the 405 nm laser line was used for excitation, and a 450/50 nm bandpass filter was used for acquisition. For the lamin B1 channel the 488 nm laser line was used for excitation, and a 520/35 nm bandpass filter was used for acquisition. For the lamin A channel the 561 nm laser line was used for excitation, and a 600/40 nm bandpass filter was used for acquisition. All the channels included a primary excitation dichroic (405/488/561/640 nm), a primary emission dichroic longpass mirror (650/ HT 660-780 nm, HR 400-640 nm), and a secondary emission dichroic shortpass mirror (568/ HT 400-550 nm, HR 620-790 nm). The three channels were acquired at a single focal plane in three sequential acquisitions in 30 randomly selected fields of view per well. >250 cells were acquired per well. For cell cycle experiments (Fig. S1C) and experiments with different cell lines (Fig. S2), a Yokogawa CV7000 high-throughput dual spinning disk microscope was used. In particular, the DAPI channel was imaged in a single plane by using the CV7000 in epifluorescence mode, with a 20X air objective (NA 0.75), a 405 nm laser as the excitation light source, a 405/488/561/640 nm excitation dichroic mirror, a 561 nm emission dichroic mirror, a 445/45 nm bandpass excitation filter, and a 2550 × 2160 pixel (5.5 MPixel) sCMOS camera with binning set to 2 (Pixel size: 650 nm). 9 randomly selected fields of view were imaged per well.

### High-throughput image analysis

The images generated by the Opera QEHS were analyzed using Columbus 2.6 (PerkinElmer). Briefly, nuclei regions of interest (ROI) were segmented using the DAPI channel, and nucleus ROI’s adjacent to the image edges were excluded from subsequent image analysis steps. The area of the nucleus ROI was measured in square microns. The mean fluorescence intensity in the lamin A and lamin B1 channels for the nucleus ROI was also measured. All the single cell values were then aggregated on a mean per well basis. Columbus per well results were exported as tab separated text files. Nuclear cross-sectional area serves as a reliable proxy for detecting differences in nuclear volume (Fig. S1A), consistent with previous reports [30,32,42,54]. Furthermore, measurements of nuclear cross-sectional areas for the same DAPI- and lamin B1-stained nuclei indicate that DAPI staining provides a generally reliable method for detecting nuclear size differences (Fig. S1B). In the case of ELYS and SEC13 knockdown, DAPI staining was preferred to lamin staining because the presence of cytoplasmic lamin puncta complicated measurements of nuclear size. For the analysis of cell cycle experiments (Fig. S1C) and experiments with different cell lines (Fig. S2), images generated by the CV7000 were imported and analyzed in Columbus 2.8.1. Nuclei were segmented as described above, and the integrated fluorescence intensity in the DAPI channels was calculated over the nucleus ROI on a per single cell basis. Analysis of cell cycle distributions obtained from single cell measurements of normalized DAPI integrated intensity was performed using R (v. 3.4.4) as previously described [114]. >650 cells per well were analyzed.

### RNAi screen statistical analysis

The statistical analysis was performed using R (v 3.3.2) and the cellHTS2 R package (v 2.36.0) [115]. Per well results were normalized on a per plate basis using the B-score method (Calculated based on the siRNA oligo library samples) in the cellHTS2 package. Normalized values for each biological replicate were then scored across the different screen plates by taking the z-score of the B-scores distribution. The biological replicates z-score values were then aggregated by calculating their mean, which is the value reported for each siRNA oligo. Putative positive hits in the RNAi screen were defined as genes that showed a z-score value of < −1.5 for at least 2 out of the 3 targeting siRNA oligos. Results for ELYS and SEC13 were validated by ordering 2 independent siRNA oligo sequences that were different from the ones used in the screen against these genes. For the validation of siRNA oligo knockdowns in different cell lines (Fig. S2E), the z-score was calculated using the mean and the standard deviation of the non-targeting negative control siRNA wells on the same plate.

### Immunofluorescence and antibodies

In general, cells were reverse transfected on poly-D-lysine coated glass coverslips (Electron Microscopy Sciences #72294-04). 48 hours post-transfection, cells were washed briefly twice with PBS and fixed with 4% PFA in PBS for 20 minutes at room temperature. All subsequent steps were performed at room temperature. After three 5-minute PBS washes, fixed cells were permeabilized with 0.5% Triton X-100 in PBS for 15 minutes at room temperature. After three 5-minute PBS washes, cells were blocked with PBS containing 0.05% Tween 20 and 5% BSA (5% PBStBSA) for 40 minutes. Cells were then incubated with primary antibodies diluted in 1% PBStBSA for four hours at room temperature or 4°C overnight. After three 5-minute washes in PBSt, cells were incubated with secondary antibodies diluted in 1% PBStBSA along with 10 μg/ml Hoechst for one hour at room temperature. After three 5-minute washes in PBSt, coverslips were mounted in Vectashield (Vector Laboratories) onto glass slides and sealed with nail polish. The following primary antibodies were used: ELYS (Santa Cruz Biotechnology, # sc-81265, mouse, 1:100), ELYS (Bethyl Laboratories, # A300-166A, rabbit, 1:50), lamin B1 (Abcam, # ab16048, rabbit, 1:1000), lamin B2 (GeneTex, # GTX628803, mouse, 1:500), lamin A/C (Santa Cruz Biotechnology, # sc-376248, mouse, 1:1000), mAb414 (BioLegend, # 902901, mouse, 1:2000), Nup133 (Santa Cruz Biotechnology, # sc-376763, mouse, 1:50), Ran (BD Transduction Laboratories, # 610341, mouse, 1:500), NTF2 (ABclonal, # A7057, rabbit, 1:200). Secondary antibodies were 1:500 dilutions of Alexa Fluor 488 and 568 anti-mouse IgG (Molecular Probes, A-11001 and A-11004) and Alexa Fluor 488 and 568 anti-rabbit IgG (Molecular Probes, A-11008 and A-11011). For FITC staining of total protein, a 10 mg/ml stock solution of FITC (Life Technologies, F1906) dissolved in DMSO was diluted to 2 μg/ml in PBS and used to stain fixed cells for one hour at room temperature.

### Microscopy and image quantification

Confocal imaging was performed on a spinning-disk confocal microscope based on an Olympus IX71 microscope stand equipped with a five line LMM5 laser launch (Spectral Applied Research) and switchable two-fiber output to facilitate imaging through either a Yokogawa CSU-X1 spinning-disk head or TIRF illuminator. Confocal images were acquired with an EM-CCD camera (ImagEM, Hamamatsu). Z-axis focus was controlled using a piezo Pi-Foc (Physik Instrumentes), and multiposition imaging was achieved using a motorized Ludl stage. Olympus objectives included UPlanFLN 40x (NA 1.30, oil) and UPlanSApo 60x (NA 1.35, oil). Image acquisition and all system components were controlled using Metamorph software. In most cases, nuclei were imaged through the largest cross-section, however in some cases the nuclear surface was imaged, for instance for measuring NPC densities. Images for measuring fluorescence intensity were acquired using the same exposure times. Total nuclear fluorescence intensities and cross-sectional nuclear areas were measured from original thresholded images using Metamorph software. Nuclear sizes were generally quantified based on images of Hoechst-stained nuclei. For NPC density measurements, images acquired on the NE surface were used to count the number of NPCs per unit area (within 5 μm^2^ regions). N/C staining intensity ratios were quantified by measuring the average staining intensity within 5 μm^2^ regions in the nucleus and cytoplasm and dividing the nuclear value by the cytoplasmic value. For publication, images were cropped and merged using ImageJ, but were otherwise unaltered. For live cell imaging, transfected cells were seeded onto chambered coverslips (Ibidi # 80826, μ-Slide 8 Well ibiTreat: #1.5 polymer coverslip). Cells were imaged 36 hours post-transfection by confocal time-lapse microscopy using objective UPlanFLN 60x (NA 0.90, air) or LCPlanFL 40× (NA 0.60, air) and a Tokai Hit stage incubator to maintain 37°C and 5% CO_2_. Images were acquired every 10-15 minutes. Time-lapse movies were analyzed and assembled using ImageJ.

### Western blots

Whole-cell lysates from tissue culture cells were prepared using SDS-PAGE sample buffer supplemented with benzonase nuclease (Sigma, E1014) and boiled for 5 min. Proteins were separated on SDS-PAGE gels (4–20% gradient) and transferred onto PVDF membrane. Membranes were blocked in Odyssey PBS Blocking Buffer (Li-Cor, 927-40000). The primary antibodies used were rabbit anti-ELYS at 1:100 (Bethyl Laboratories, # A300-166A), mouse anti-CRM1/XPO1 at 1:100 (Santa Cruz Biotechnology, # sc-74454), and DM1A mouse anti-α-tubulin at 1:2000 (Santa Cruz Biotechnology, # sc-32293). The secondary antibodies were IRDye-680RD-conjugated anti-mouse-IgG (Li-Cor 925-68070) and IRDye-800CW-conjugated anti-rabbit-IgG (Li-Cor 926-32211) used at 1:20,000. Blots were scanned on a Li-Cor Odyssey CLx instrument, and band quantification was performed with ImageStudio, normalizing to the tubulin signal.

### Statistics

Where indicated, nuclear area and intensity measurements were normalized to controls. Averaging and statistical analysis were performed for independently repeated experiments. Two-tailed Student’s t-tests assuming equal variances were performed in Excel (Microsoft) to evaluate statistical significance. The p-values, number of independent experiments, number of nuclei quantified, and error bars are denoted in the figure legends.

## ACKNOWLEDGEMENTS

Research in the Levy lab is supported by funding from the National Institutes of Health/National Institute of General Medical Sciences (R01GM113028) and the American Cancer Society (RSG-15-035-01-DDC). Research in the Misteli lab and HiTIF is supported by funding from the Intramural Research Program of the National Institutes of Health (NIH), National Cancer Institute, and Center for Cancer Research. The authors declare no competing financial interests.

## AUTHOR CONTRIBUTIONS

Conceptualization, P.J., A.C.S., G.P., T.M., D.L.L.; Methodology, A.C.S., G.P., T.M.; Formal Analysis, G.P.; Investigation, P.J., A.C.S., G.P.; Writing – Original Draft, P.J., D.L.L.; Writing – Review & Editing, P.J., A.C.S., G.P., T.M., D.L.L.; Funding Acquisition, T.M., D.L.L.; Supervision, T.M., D.L.L.

## SUPPLEMENTAL FIGURE LEGENDS

**Figure S1:**
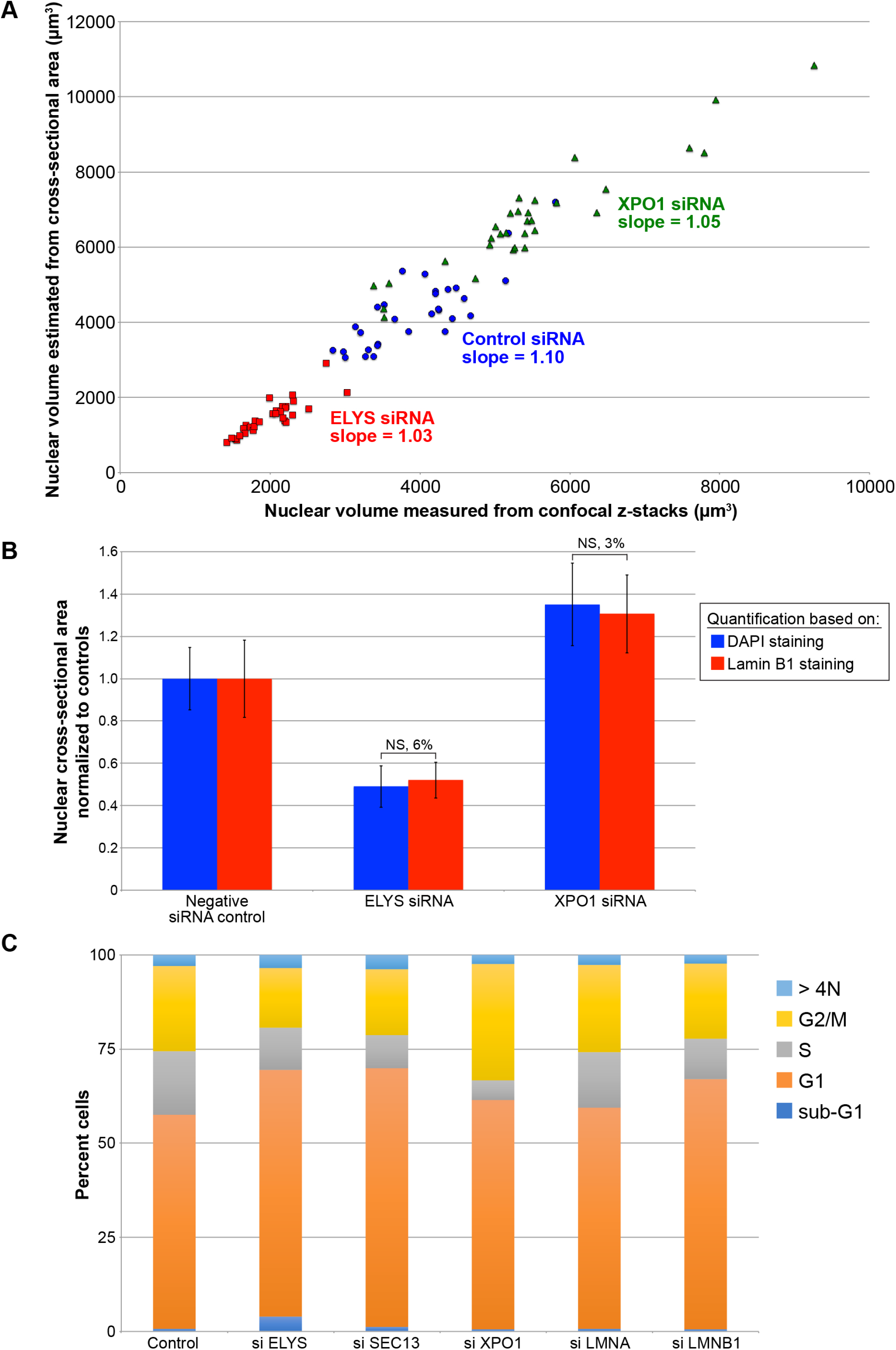
Validation of nuclear size quantification methods. (**A**) MCF-10AT1k.cl2 cells were transfected with control siRNA, ELYS siRNA, or XPO1 siRNA and stained with DAPI and an antibody against lamin B1. Confocal z-stacks were acquired with a z-slice thickness of 1 μm and a total of 21 z-slices. Nuclear volume was quantified from the DAPI confocal z-stacks and plotted on the x-axis. For the same DAPI-stained nuclei, nuclear volume was estimated from the maximum cross-sectional nuclear area assuming a spherical nucleus and plotted on the y-axis. Data from three biological replicates are shown. For each experimental condition, 30 nuclei were quantified, linear regression analysis was performed, and the slope of the fitted line is indicated. A t-test calculated by comparing all measured nuclear volumes to estimated nuclear volumes returned a non-significant p-value of 0.25. (**B**) Maximum nuclear cross-sectional areas were quantified from the confocal z-stacks described in (**A**) for both the DAPI and lamin B1 channels. For each experimental condition, three biological replicates were performed and 30 nuclei were quantified. For a given staining method, nuclear areas were normalized to the negative siRNA control. The percent difference between the DAPI and lamin B1 staining methods is indicated above each pair of bars. (**C**) MCF-10AT1k.cl2 cells were transfected with control siRNA or with siRNA against the indicated genes. Cells were stained with DAPI and quantification of DNA staining intensity was used to estimate the fraction of cells in various stages of the cell cycle by high-throughput imaging as previously described [114] (see Methods). The stacked bars represent the means of the fractions for each cell cycle phase calculated over 3 biological replicates. NS, not significant; error bars represent SD.

**Figure S2:**
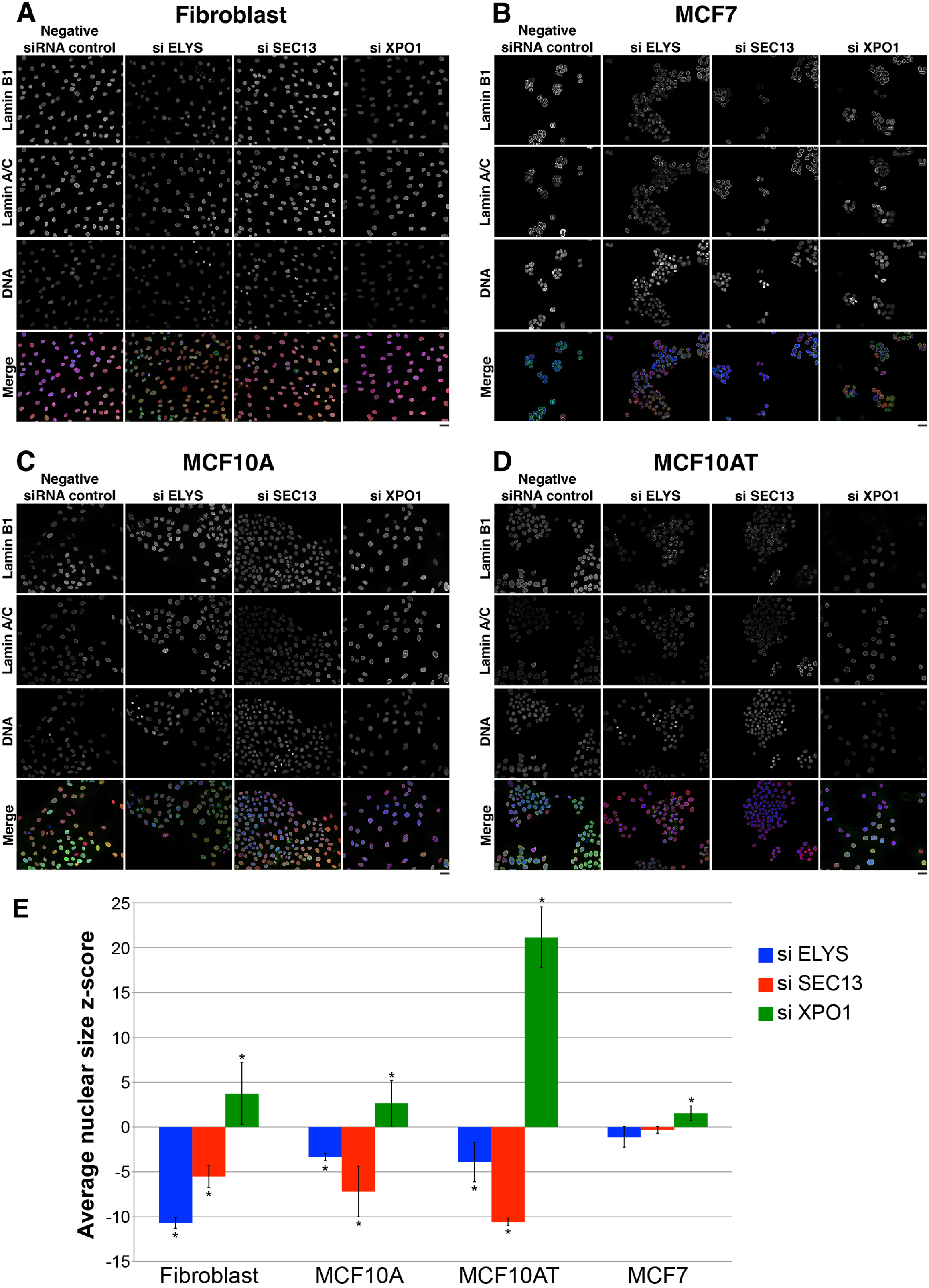
ELYS, SEC13, and XPO1 knockdown in different cell lines. The indicated cell lines were transfected with a negative siRNA control or siRNA against ELYS, SEC13, or XPO1. Cells were stained with DAPI and with antibodies against lamin B1 and lamin A/C. (**A**) Representative images for hTERT immortalized CRL-1474 cells, roughly normal human skin fibroblasts. (**B**) Representative images for MCF7 cells, breast adenocarcinoma cells. (**C**) Representative images for MCF-10A cells, roughly normal breast epithelial cells. (**D**) Representative images for MCF-10AT1k.cl2 cells, breast epithelial cells with atypical hyperplasia. (**E**) Nuclear cross-sectional areas were quantified and z-scores were calculated as described in Methods. For each cell line and condition, >500 cells were quantified and the average z-score and SD from three biological replicates are plotted. *, absolute value of the z-score is greater than 1.5; scale bars are 25 μm.

**Figure S3:**
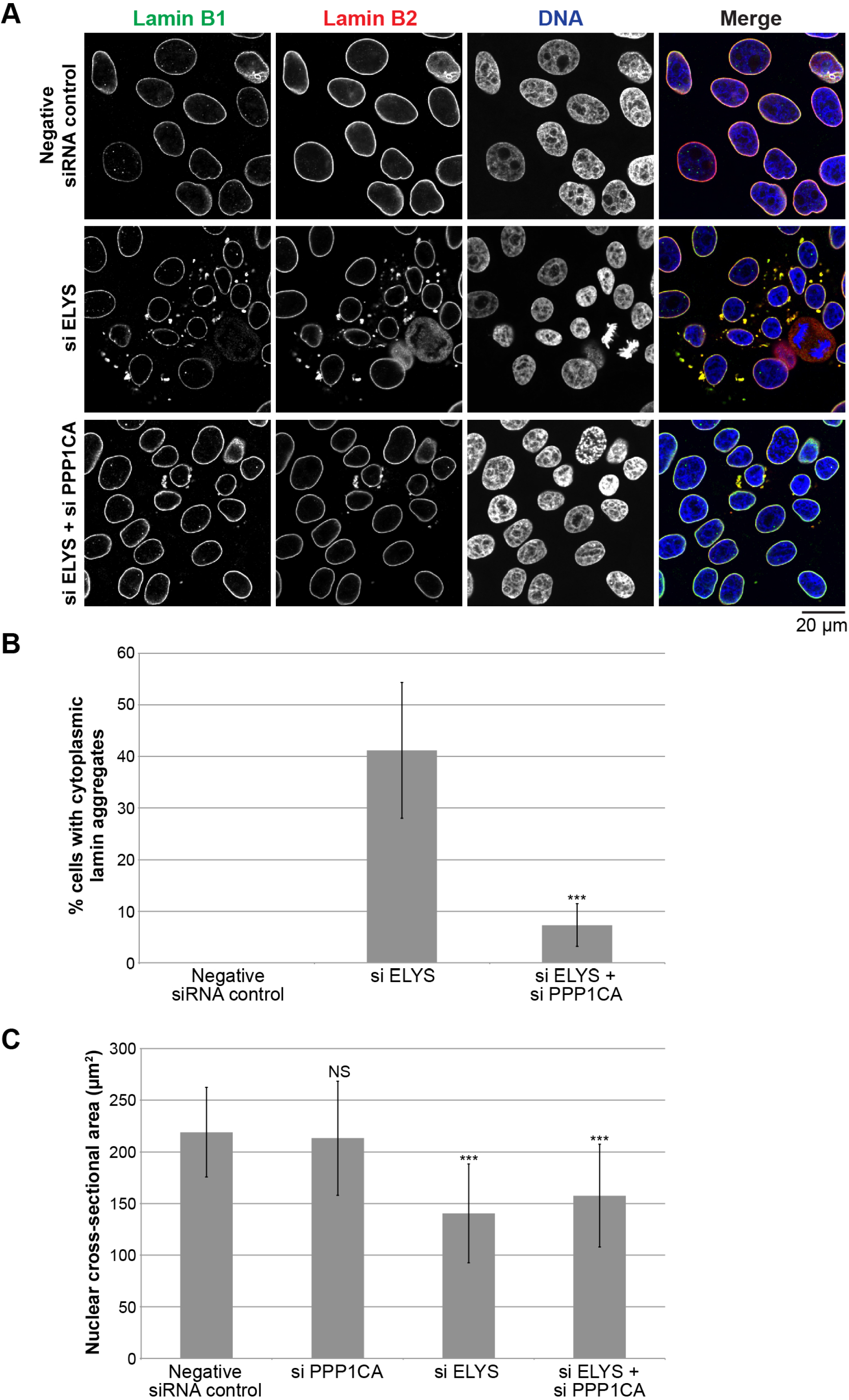
Protein phosphatase PP1α knockdown rescues the lamin aggregation phenotype in ELYS knockdown cells. (**A**) MCF-10AT1k.cl2 cells were transfected with control siRNA, ELYS siRNA, or ELYS siRNA plus PPP1CA siRNA (protein phosphatase 1 catalytic subunit alpha) and stained with antibodies against lamin B1 and B2. Representative images are shown. (**B**) Cells with and without cytoplasmic lamin B aggregates were counted and the percentage of cells with aggregates was calculated. For each experiment, 28-120 cells were examined per condition. Data from one representative experiment of two are shown. (**C**) For each experiment, nuclear cross-sectional areas were quantified for 70-184 nuclei per condition (117 nuclei on average) and averaged. Data from one representative experiment of two are shown. ***, p<0.005; NS, not significant; error bars represent SD.

**Figure S4:**
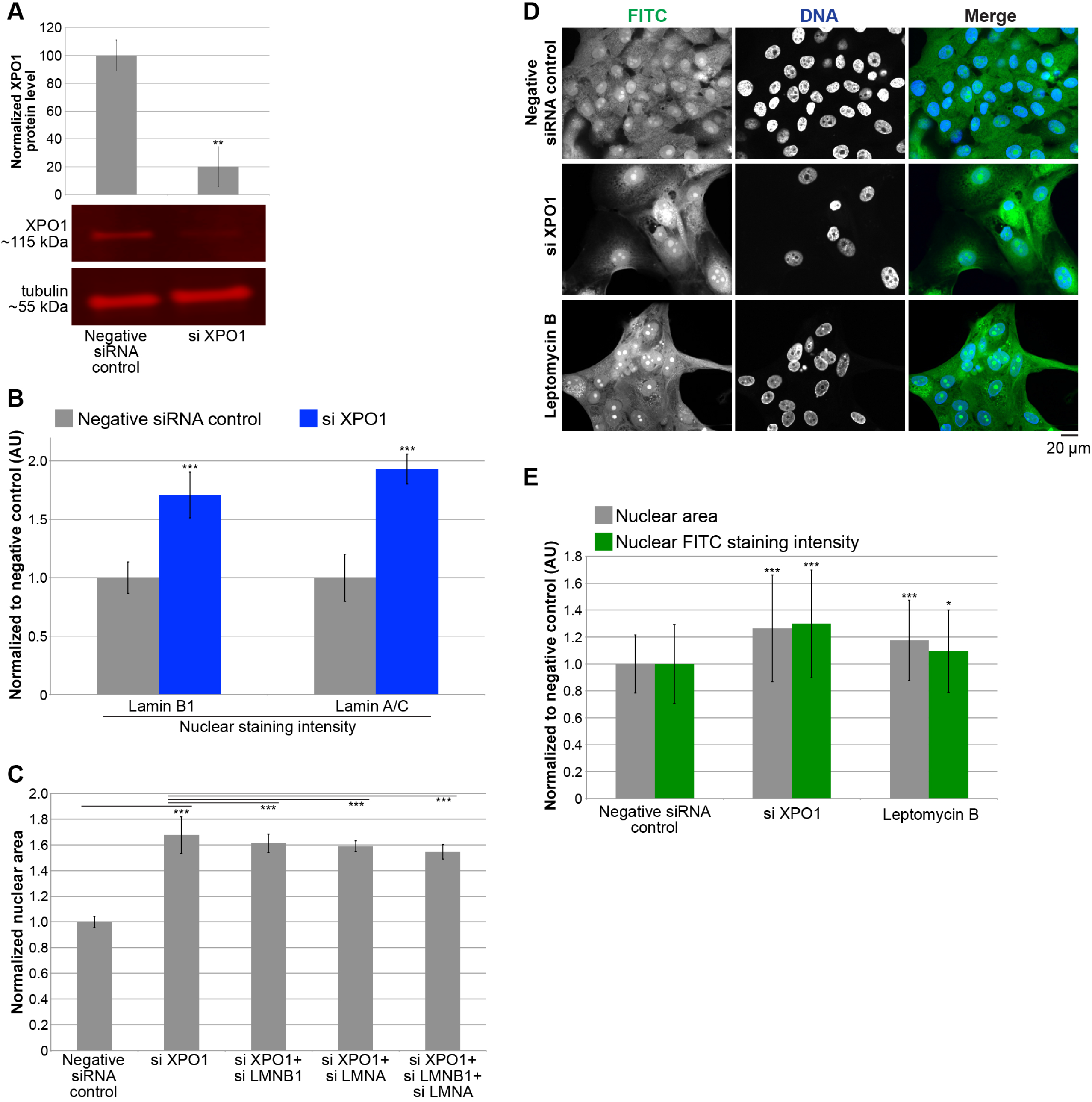
Reducing nuclear export increases nuclear size. Representative images for XPO1 siRNA are shown in Figure S2. (**A**) Cell lysates from control and XPO1 siRNA transfected cells were analyzed by western blot and probed for XPO1 and tubulin. One representative western blot is shown. XPO1 band intensity was normalized to tubulin. Quantification from two biological replicates is shown. (**B**) MCF-10AT1k.cl2 cells were transfected with control or XPO1 siRNA. Lamin B1 and lamin A/C nuclear staining intensities were quantified based on experiments using two different XPO1 siRNA sequences and two biological replicates. For each experiment, 108-865 nuclei were quantified per condition (416 nuclei on average). Average and SD values based on averages from these four experiments are plotted, normalized to the negative siRNA control. (**C**) MCF-10AT1k.cl2 cells were transfected with the indicated siRNAs and nuclear cross-sectional areas were quantified. For each experiment, 203-604 nuclei were quantified per condition (403 nuclei on average). Average and SD values based on averages from four biological replicates are plotted, normalized to the negative siRNA control. (**D**) MCF-10AT1k.cl2 cells were transfected with control or XPO1 siRNA. For the bottom row of images, cells transfected with control siRNA were treated with 20 ng/ml leptomycin B for 24 hours. Cells were fixed and stained with 2 μg/ml FITC. Representative images are shown. (**E**) For each experiment, nuclear cross-sectional areas and nuclear FITC staining intensities were quantified for 105-620 nuclei per condition (267 nuclei on average) and normalized to the negative control. Data from one representative experiment of two are shown. ***, p<0.005; **, p<0.01; *, p<0.05; error bars represent SD.

## MOVIE LEGENDS

**Movie 1**: MCF-10AT1k.cl2 cells were co-transfected with ELYS siRNA and with plasmids expressing mCherry-lamin B2 and H2B-GFP. Live imaging was performed at 10-minute intervals, for a total of 23 hours. Some out of focus frames were removed. This movie shows cytoplasmic lamin B2 puncta that appear during interphase and that are initially small but appear to coalesce into larger puncta over time. Lamin B2 puncta display dynamic movements within the cell, and the disappearance of puncta can also be observed. It is worth noting the transfected cell to the left that exhibits no obvious cytoplasmic lamin B2 puncta and the largest nucleus in the field of view.

**Movie 2**: MCF-10AT1k.cl2 cells were co-transfected with ELYS siRNA and with plasmids expressing mCherry-lamin B2 and H2B-GFP. Live imaging was performed at 15-minute intervals, for a total of 9 hours. A cell without obvious cytoplasmic lamin B2 puncta divides to produce daughter cells that exhibit lamin B2 puncta in interphase. There is also a cell in the upper left corner with abundant cytoplasmic lamin B2 foci and no nuclear localized lamin B2.

**Movie 3**: MCF-10AT1k.cl2 cells were co-transfected with ELYS siRNA and with plasmids expressing mCherry-lamin B2 and H2B-GFP. Live imaging was performed at 15-minute intervals, for a total of 14.5 hours. Some out of focus frames were removed. A cell with cytoplasmic lamin B2 puncta divides to produce daughter cells that initially exhibit lamin B2 puncta in interphase that appear to dissolve.

**Movie 4**: MCF-10AT1k.cl2 cells were co-transfected with ELYS siRNA and with plasmids expressing mCherry-lamin B2 and H2B-GFP. Live imaging was performed at 15-minute intervals, for a total of 19 hours. Some out of focus frames were removed. This movie shows dynamic movements of cytoplasmic lamin B2 puncta.

**Movie 5**: MCF-10AT1k.cl2 cells were co-transfected with ELYS siRNA and with plasmids expressing mCherry-lamin B2 and H2B-GFP. Live imaging was performed at 15-minute intervals, for a total of 4 hours. Some out of focus frames were removed. A cell with cytoplasmic lamin B2 puncta divides to produce daughter cells with lamin B2 puncta in interphase.

**Movie 6**: MCF-10AT1k.cl2 cells were co-transfected with ELYS siRNA and with plasmids expressing mCherry-lamin B2 and H2B-GFP. Live imaging was performed at 15-minute intervals, for a total of 2 hours. A representative cell is shown in which ectopically expressed mCherry-lamin B2 exists exclusively in cytoplasmic foci with none localized to the nuclear envelope.

**Table S1:** Gene knockdowns that reduced nuclear size in MCF-10AT1k.cl2 cells.

| **Gene** | **max. z-score** | **SEM** | **Gene** | **median z-score** | **SEM** |
| --- | --- | --- | --- | --- | --- |
| SMARCC2 | -3.48 | 0.8355 | SEC13 | -2.65 | 0.2245 |
| RAI1 | -3.26 | 0.2055 | SHPRH | -2.53 | 0.3855 |
| ITGAX | -3.23 | 0.3475 | FZR1 | -2.37 | 0.3525 |
| RAC1 | -3.07 | 0.1270 | SMARCC2 | -2.29 | 0.2225 |
| SEC13 | -2.97 | 0.2000 | RAC1 | -1.98 | 0.2825 |
| SHPRH | -2.70 | 0.8340 | AHCTF1 | -1.97 | 0.1560 |
| AHCTF1 | -2.47 | 0.2415 | NCAPD2 | -1.94 | 0.3815 |
| FZR1 | -2.42 | 0.0930 | ATXN7L3 | -1.82 | 0.4360 |
| ATXN7L3 | -2.32 | 0.0665 | ITGAX | -1.75 | 0.1370 |
| CMTM5 | -2.24 | 0.3475 | SUN3 | -1.75 | 0.3610 |
| HAT1 | -2.13 | 0.1345 | WDR61 | -1.67 | 0.1330 |
| NCAPD2 | -1.98 | 0.1145 | RAI1 | -1.63 | 0.4070 |
| TADA3 | -1.97 | 0.2560 | UBE2E1 | -1.62 | 0.0735 |
| SUPT16H | -1.90 | 0.2460 | CREBBP | -1.61 | 0.3660 |
| WDR61 | -1.88 | 0.0195 | BMI1 | -1.58 | 0.5715 |
| BMI1 | -1.82 | 0.2505 | HAT1 | -1.57 | 0.2110 |
| SUN3 | -1.76 | 0.2490 | CMTM5 | -1.55 | 0.1830 |
| CREBBP | -1.70 | 0.7185 | TADA3 | -1.54 | 0.2760 |
| UBE2E1 | -1.67 | 0.0090 | SUPT16H | -1.51 | 0.2575 |

Z-scores were calculated for three different siRNA oligo sequences for each gene and two biological replicates.

**Table S2:** Function of hits that reduced nuclear size in MCF-10AT1k.cl2 cells.

| **Gene** | **Function** |
| --- | --- |
| *SMARCC2* | Component of the SWI/SNF chromatin remodeling complex |
| *RAI1* | Circadian clock transcriptional regulator |
| *ITGAX* | Receptor for fibrinogen |
| *RAC1* | Small GTPase associated with the plasma membrane |
| *SEC13* | Component of the Nuclear Pore Complex |
| *SHPRH* | E3 Ubiquitin Ligase |
| *AHCTF1* | Factor required for Nuclear Pore Assembly |
| *FZR1* | Component of the Anaphase Promoting Complex/Cyclosome |
| *ATXN7L3* | Component of SAGA (histone acetyltransferase complex) |
| *CMTM5* | Transmembrane domain containing protein |
| *HAT1* | Lysine Acetyltransferase |
| *NCAPD2* | Component of the Condensin Complex |
| *TADA3* | Component of the PCAF complex |
| *SUPT16H* | Component of the FACT complex |
| *WDR61* | Component of the PAF1 complex |
| *BMI1* | Component of the PRC-1 like complex |
| *SUN3* | Component of the LINC complex |
| *CREBBP* | Lysine Acetyltransferase |
| *UBE2E1* | E2 ubiquitin conjugating enzyme |

## Abbreviations

NE: nuclear envelope
NPC: nuclear pore complex
ER: endoplasmic reticulum
Nup: nucleoporin
FG-Nup: phenylalanine-glycine repeat nucleoporin

